# Neighbourhood effect of weeds on wheat root endospheric mycobiota

**DOI:** 10.1101/2022.09.02.506399

**Authors:** Jie Hu, Claire Ricono, Paola Fournier, Samuel Mondy, Philippe Vandenkoornhuyse, Cendrine Mony

**Author notes:** The authors have contributed equally (Jie Hu, Claire Ricono).

## Abstract

1. Microorganisms associated with plants provide essential functions to their hosts, and therefore affect ecosystem productivity. Agricultural intensification has modified microbial diversity in the soil reservoir and may affect plant microbial recruitment. Weeds develop spontaneously in crop fields, and could influence microorganisms associated with crop plants through a neighbourhood effect. We explore the effect of weed species on crop plant microbiota as potentially auxiliary plants that affect agricultural productivity.
2. We combined field and controlled laboratory studies to analyse the neighbourhood effect of weeds on wheat root endospheric mycobiota and growth. First, we analysed the effect of weed species diversity and identity recorded in the neighbourhood of individual wheat plants on soil and wheat root mycobiota in the field. Second, we used a plant-matrix design in laboratory conditions to test the effect of weed identity (9 weed treatments) and their ability to transmit root mycobiota to wheat roots, and the resulting impact on wheat growth.
3. In contrast to soil mycobiota, we demonstrated that wheat root endospheric mycobiota was influenced by the diversity and identity of weeds developing in their 1 m^2^ neighbourhood. Wheat root endospheric microbiota strongly differs in terms of richness and composition depending on the neighbouring weed plant species. Weed species transmitted from 13% to 74% of their root microbiota to wheat roots depending on weed identity in controlled conditions.
4. ***Synthesis.*** Weed neighbours modified wheat plant performance, possibly as a result of competitive interactions and changes in microbiota. Our findings suggest that crop root mycobiota was variable and was modulated by their weed neighbourhood. Synergistic effects between mycobiota of crops and weeds could therefore contribute to soil biodiversity and sustainable agriculture.

## 1 Introduction

Plants harbour diverse microorganisms in and on their tissues, forming their associated microbiota (Berg et al., 2016). Plant associated microbiota fulfil essential functions for plant nutrition (Hardoim et al., 2015), plant protection against abiotic stress (Lenoir et al., 2016) and plant immune system (Hacquard et al., 2017). Maintaining or even engineering plant associated microbiota can therefore help boost plant yields in a sustainable way (Busby et al., 2016). However, today’s intensive agricultural systems have led to a microbial diversity crisis, caused, for example, by agrochemical application, mechanical management, crop rotation reduction and monospecific plant assemblages leading to global loss of biodiversity in agroecosystems (Creamer et al., 2016; Hartman et al., 2018). This microbial diversity crisis may affect plant fitness and productivity through detrimental recruitment of its microbiota, especially that of plant endophytic microbiota.

Plants recruit their microbiota in the local soil reservoir (Vandenkoornhuyse et al., 2015), and recruitment is in part deterministic (Guo et al., 2021; Wippel et al., 2021). Environmental factors and the dispersal capacity of microorganisms shape the microbial reservoir in ecosystems (Fierer, 2017; Martiny et al., 2006). Plants recruit microorganisms in soil reservoir through a filtering process related to plant morphological, chemical and biological traits such as root type (Saleem et al., 2018), root exudate profile (Haichar et al., 2008) and plant immunity (Dodds & Rathjen, 2010). In addition, some rewarding processes that promote root colonisation by specific fungi that are the most cooperative for the plants (Kiers et al., 2011). These active and passive filtering processes have led to a certain level of host preference which can be observed both at the species and genotypic level. For instance, (Xiong et al., 2021) showed that crop identity (maize, rice or wheat) mainly determined microbiota recruitment rather than the field location or fertilisation management. Distinct root-associated microbial communities have been reported in phylogenetically distant plants, including maize, sorghum and wheat (Bouffaud et al., 2014), among close plant relatives such as *Arabidopsis* and *Cardamine hirsuta* (Schlaeppi et al., 2014), and even different cultivars such as rice (Andreo-Jimenez et al., 2019). Interactions between individual plants and their associated microorganisms are well described (Hardoim et al., 2015). However, *in situ* plant-microbe interactions occur in a more complex biotic context where monospecific plant assemblages are the exception, and multispecies assemblages or spontaneous flora developing together with crop plants are the norm. Consequently, little is known about how plant-plant interactions in multispecies assemblages affect plant-microbe interactions, particularly their associated microbiota.

Recent works suggest a plant neighbourhood effect on a focal plant endopheric microbiota in grassland mesocosms (Bittebiere et al., 2020; Mony et al., 2020). The identity of plants growing within a few centimetres of the focal plant were shown to affect the richness and composition of the root endospheric mycobiota associated with *Brachypodium pinnatum*. This neighbourhood effect could be caused indirectly by root exudate production that can modify local soil microbiota (Saunders et al., 2010) via favouring or rejecting specific microorganisms. (Steinauer et al., 2016) reported that specific mixtures of root exudates can modify soil microbial composition, and the chemical class of root exudates accurately predicted changes in microbial composition and diversity (Gu et al., 2020). Neighbouring plants can also directly transfer part of their microbiota to focal plants (Mony et al., 2021). This transmission can be achieved through root contact or small-scale dispersal (Enkhtuya et al., 2005; Smith & Read, 2008). But how and to what extent the identity and diversity of neighbouring plants and their associated microbiota can affect the microbiota and its consequences on the fitness of the plants developing in this neighbourhood need to be investigated more thoroughly.

In agrosystems, cultivated crop plants are usually spontaneously surrounded by weed plants. Agricultural fields harbour a large seedstock of weed plants that contribute to a varied population of neighbouring plants for crops, especially under organic management (Armengot et al., 2013). Weeds are thus likely to influence the microbiota of crop plants through direct contact or indirect modification of the soil microbial reservoir. Weed species vary in their ability to recruit microbiota for themselves and may also shape the diversity and abundance of microorganisms in the soil. Furthermore, weeds may influence the productivity of crop plants through changes in their functional microbiota. For instance, experimental removal of particular weed species in fields, which resulted in modifications in the AMF composition associated with crops, led to a reduction of their beneficial effects on plant productivity in the field (Feldmann & Boyle, 1999; Kabir & Koide, 2000). The potential positive role of weeds led to a debate with farmers that endorsed the paradigm that weed species compete with crops for resources, reduce crop yields and have to be removed, in addition to their emerging resistance to herbicides (Llewellyn et al., 2004). Moreover, it has been proposed that we need to better understand the relationship between weeds and crops in agrosystem functioning and agricultural management (Carlos et al., 2014). In agricultural context, the importance of weeds for the microbial compartment has been overlooked up to now.

In this study, we analysed how the mycobiota associated with a crop plant can be influenced by weeds. We focused on the influence of weed diversity and identity on soil mycobiota and on wheat root endospheric mycobiota in fields under organic management. First, we analysed the effect of composition and richness of neighbouring weeds on soil mycobiota and wheat root endospheric mycobiota in a set of organic fields by sampling individual wheat plants surrounded by different local weed plant neighbourhoods. Second, we conducted an experiment in controlled laboratory conditions on nine weed species selected based on field data to analyse how weed root endospheric mycobiota affects wheat performance via the transmission of weed root mycobiota to crop roots.

We hypothesised that (1) in field conditions: (i) weed species diversity shapes the composition and enriches the fungi of mycobiota in the soil and in that associated with crop roots; (ii) the identity of weed species in the neighbourhood plays a specific role in these relationships; (2) in controlled conditions: (i) the composition and diversity of the mycobiota associated with the roots of weed species affect their ability to transmit mycobiota to individual wheat plants; (ii) when the mycobiota of weed plant species is transmitted to wheat plants, there is a change in the composition and diversity of the wheat root mycobiota; (3) transmitted mycobiota compensate for focal wheat plant growth, especially when there is no or only a limited microbial reservoir.

## 2 Method and materials

### 2.1 Field study

We selected 15 organic winter wheat fields in the Long-Term Socio-ecological Research (LTSER) site “*Zone Atelier Armorique*”, located in north-western France (48°06’43”N 1°40’27”W). The 15 fields are located in a bocage area, in a mosaic of agricultural fields partly surrounded by hedgerows, and characterised by mixed crop-livestock farming. The fields were managed via using tillage and mechanical deweeding and no plant protection products or pesticides were used for field and hedgerow management (Ricono et al., 2022). In each field, we selected four sampling points located at least 10 metres from the edge of the field to avoid edge effects.

At each sampling point, we collected soil and wheat roots when individual wheat plants were at the reproductive stage. At each location where the samples of soil and wheat were collected, we performed floristic surveys in 1 × 1 m quadrats to identify the floristic neighbourhood of each individual wheat plant. In each quadrat, we visually estimated the percentage cover of each weed species. Then in each plant neighbourhood, we identified the composition and abundance of the weed community.

### 2.2 Controlled experiment

We analysed the influence of weed neighbour species on wheat root endospheric mycobiota in a controlled experiment using a plant-matrix design where individual wheat plants were planted in a matrix of four individuals of the same weed species (Figure S1, Table S1). We used the winter wheat variety Attlass and focused on weed species that were (i) frequent in wheat fields, (ii) that had sufficient root biomass to enable molecular analysis, (iii) were representative of different plant families; and (iv) of which wild seeds were available without domestication by breeders. We selected nine weed species as a subsample of all the weed species found in the field. Ten replicates of each treatment were performed using a neighbour of a single weed species (i.e. 9 treatments (i) *Galium aparine* (ii) *Lamium purpureum* (iii) *Matricaria sp.* (iv) *Papaver rhoeas* (v) *Poa annua* (vi) *Poa trivialis* (vii) *Trifolium repens* (viii) *Veronica persica* and (ix) *Vicia sativa*), with two additional control treatments (i.e. a single wheat plant grown alone, and an individual wheat plant surrounded by four sterile wheat plants). For each replicate, the roots of the focal plant and of the neighbouring individual plants were sampled to characterise the associated endospheric mycobiota. Wheat and weed aboveground dry biomass were also measured as a proxy of wheat fitness. More details about experimental design were included in supplementary materials.

### 2.3 Soil and root mycobiota analysis

#### Sample preparation

From the composite soil sample from each plot, a homogenised aliquot of soil was sieved to 4 mm and 50 g of soil were sent to the Genosol platform for lyophilization or stored at −40 °C before DNA extraction. From each individual plant sampled, 80 mg of roots were washed in tap water for 5 mins, then placed in a 20-mL sterile polypropylene tube with a 5‰ Triton X100 solution for 10 mins. Finally, the roots were thoroughly rinsed with sterile 18mΩ purified water. Small pieces of root (< 1 cm) were sampled randomly from different parts of the root system of each individual wheat plant, and 80-mg aliquots of roots were stored in 1.5 mL Eppendorfs^®^ tubes at −20 °C before DNA extraction along with samples taken from subsequent controlled experiments.

#### DNA extraction, 18S rRNA amplicon sequencing and bioinformatics

DNA from soil samples was extracted at the GenoSol Platform. DNA was extracted from the sample roots of all the weed and wheat plants from both the field study and controlled experiments at the Gentyane platform. We used 18S rRNA to analyse the root fungal endospheric mycobiota of the wheat and weed plants. All PCR products were purified with AMpureXP magnetic beads (Agencourt^®^) using an automated liquid platform (Bravo-Agilent^®^) and quantified (Quant-iT PicoGreenTM dsDNA Assay Kit) to allow DNA normalization at the same concentration, and a second round of PCR, purification, quantification, library construction and sequencing step were performed at the ‘EcogenO’ platform.

Data trimming consisted of removing primer and degenerated base sequences (Cutadapt). Trimmed sequences were then analysed using the FROGS pipeline (Escudié et al., 2018). We used the FROGS standard pre-process to process the sequence data. This pipeline uses SWARM for cluster formation. The PhymycoDB database (Mahé et al., 2014) was used for fungal 18S rRNA gene sequence affiliation. Based on the rarefaction curves drawn for each dataset (Figure S2), contingency matrices were normalized to 21,743 reads for soil mycobiota, 14,530 reads for wheat root endospheric mycobiota for the field study, and 4,203 for wheat and weed root endospheric mycobiota for the controlled experiment. Samples under these thresholds were removed. More details about DNA extraction, 18S rRNA amplicon sequencing and bioinformatics were included in supplementary materials.

#### Mycobiota parameter calculation

In both studies, the number of sequences per sample made it possible to describe the root endospheric fungal assembly in sufficient depth (curve slopes asymptotically close to 0). A total of 60 soil mycobiota samples were analysed, 60 wheat root endospheric mycobiota samples in the field (15 × 4 sampling points) in the field study; and 93 wheat and 84 weed root mycobiota samples were analysed in the controlled experiment (7 wheat root samples and 6 weed root samples were discarded due to low quality or quantity of DNA or PCR products). All statistical analyses were performed on these normalized contingency matrices.

We calculated the diversity of the soil and wheat root endospheric fungal communities based on the normalized contingency matrices in the field study, including diversity (hereafter sequence cluster richness) and Pielou’s evenness index. These metrics were calculated for the ‘all fungi’ and for the five most frequently represented phyla (Ascomycota, Basidiomycota, Chytridiomycota, Glomeromycota and Zygomycota) in the soil mycobiota and in the wheat root endospheric mycobiota.

The diversity of the wheat and weed root endospheric fungal community was also calculated based on the normalized contingency matrices in the controlled experiment, including sequence cluster richness, Pielou’s evenness index, the number of shared sequence clusters using the R vegan package and the percentage of shared sequence clusters. The percentage of sequence clusters shared by wheat and weeds in the pot experiment was calculated as the ratio of the number of sequence clusters shared by weeds and wheat to the number of sequence clusters of the weeds alone.

### 2.4 Statistical analyses

#### Field survey

A Venn diagram was drawn using the R package VennDiagram to detect the shared and single sequence clusters in the soil mycobiota and wheat root endospheric mycobiota in the organic fields. Two co-inertia multivariate analyses (Doledec & Chessel, 1994) were then performed to determine if the composition of the weed neighbourhood was related to the soil mycobiota or to the wheat root endospheric mycobiota. For this purpose, only sequence clusters and plant species that were found in at least 3% of the samples were used. The significance of the co-inertia was tested using the Monte-Carlo permutation test with the “randtest” function in the “ADE4” package. In addition, the effects of weed richness on the composition of the soil and wheat root endospheric mycobiota with PERMANOVA were tested using the adonis function of the vegan package in R. The effects of weed richness on sequence cluster richness of ‘all fungi’ and of each phylum in the soil mycobiota and wheat root mycobiota in the field sites were tested using a mixed model with negative binomial distributions in the R package “lme4” (Bates et al., 2015). The field site was used as a random factor to control for data dependency (4 samples per field site). The normality and homoscedasticity of the model were checked using a graphical representation of the residuals. The marginal (R^2^m) and conditional (R^2^c) values of R^2^ were calculated for all models. These R^2^ corresponded to the variance explained by the fixed effects and the addition of fixed and random effects, respectively. A Tukey post-hoc test was used for group comparisons of sequence cluster richness and evenness of soil mycobiota diversity and wheat root endospheric mycobiota in the field.

#### Controlled experiment

We used principal coordinate analysis (PCoA) to identify the composition of the wheat and weed root endospheric mycobiota communities in combined and separate analyses. The least significant difference was also used via the LSD.test in the agricolae package to compare each weed species along the first and second principal components of the weed and wheat root endospheric mycobiota. We also identified the sequence clusters that were enriched or depleted in wheat root endospheric mycobiota depending on the neighbourhood species. For this purpose, we conducted log2foldchange analysis using R package DESeq2 (Love et al., 2014) to compare each sequence cluster in the root mycobiota of wheat with weeds as neighbours to each sequence cluster in the root mycobiota of wheat in the control treatment without any weed neighbours. After calculation, the sequence clusters whose abundance of log2foldchange was higher than 0.6 or lower than −0.6 and with a significant P value were kept to count the amount of changed (both enriched and reduced sequence clusters in each treatment.

The effect of weed-mediated change in root endospheric mycobiota on wheat performance was assessed through aboveground biomass. The effect of weed identity on wheat aboveground dry biomass was tested along with the effect of wheat root endospheric mycobiota diversity (i.e. “all fungi” sequence cluster richness, sequence cluster richness in the phyla Ascomycota, Basidiomycota, Chytridiomycota, Glomeromycota and Zygomycota) on wheat aboveground dry biomass. In both cases, generalised linear models were used. Significance was tested using a Type II ANOVA after checking for normal distribution of residuals. Linear models were used to detect the effects of weed identity on wheat and weed root endospheric mycobiota sequence cluster richness, on the number of shared sequence clusters, the percentage of shared sequence clusters, the weight of wheat and weed aboveground plant biomass in the controlled experiment. A Tukey post-hoc test was used for group comparisons of sequence cluster richness of weed and wheat root endospheric mycobiota, the number and percentage of sequence clusters shared by weeds and wheat. All statistical analyses were performed using R software (R Development Core Team, 2013) version 4.0.0.

## 3 Results

### 3.1 Effects of weed neighbourhood on wheat root mycobiota in the field study

A co-inertia analysis showed that, except for Glomeromycota, soil mycobiota was not influenced by floristic composition in the neighbourhood (Table 1). Floristic richness did not affect the composition (Table 2), the sequence cluster richness or the evenness of “all fungi” and each phylum of the soil mycobiota (Table S2), indicating a very limited legacy effect of weed species on the soil microbial reservoir. However, we found a significant relationship between floristic composition in the neighbourhood of wheat individuals and the endospheric mycobiota associated with wheat roots, particularly for “all fungi” and phylum Zygomycota (Table 1). Floristic richness and evenness significantly affected the composition of wheat root endospheric mycobiota (Table 2). Floristic richness increased wheat root endospheric mycobiota sequence cluster richness for the whole fungi, in the phyla Ascomycota, Glomeromycota and Zygomycota, and floristic richness increased wheat root endospheric mycobiota sequence cluster evenness in the phylum Basidiomycota (Table S2).

**Table 1.** Co-inertia analysis between floristic composition and soil mycobiota, and between floristic composition and root endospheric mycobiota of wheat in the field study. The RV coefficients obtained by co-inertia analysis between the same paired data sets highlight the relationship between the floristic species abundance and mycobiota sequence cluster relative abundance of soil or wheat root endosphere. Total inertia of co-inertia is related to the explained variance supported by its 2 first axes. P values were calculated using a Monte-Carlo test based on 999 permutations. Significant results (P < 0.05) are highlighted in bold, and marginal significant results (0.05 < P < 0.10) are highlighted in bold and italics.

|  | RV | Total inertia: Axis 1&2 (%) | P |
| --- | --- | --- | --- |
| **Soil mycobiota** |  |  |  |
| All fungi | 0.55 | 19.9 | 0.261 |
| Ascomycota | 0.49 | 22.9 | 0.227 |
| Basidiomycota | 0.43 | 30.4 | 0.584 |
| Chytridiomycota | 0.37 | 30.6 | 0.569 |
| Glomeromycota | 0.34 | 49.6 | **0.012** |
| Zygomycota | 0.36 | 31.1 | 0.547 |
| **Wheat root endospheric mycobiota** |  |  |  |
| All fungi | 0.56 | 31.5 | **0.020** |
| Ascomycota | 0.49 | 30.4 | 0.104 |
| Basidiomycota | 0.46 | 31.9 | **0.082** |
| Chytridiomycota | 0.34 | 33.1 | 0.566 |
| Glomeromycota | 0.29 | 49.8 | 0.169 |
| Zygomycota | 0.43 | 48.9 | **0.002** |

**Table 2.** Effect of floristic diversity on soil mycobiota and wheat root endospheric mycobiota composition in the field study. Floristic diversity is indicated as floristic richness and evenness. Effects were tested via a PERMANOVA analysis. Significant results (P < 0.05) are highlighted in bold, and marginal significant results (0.05 < P < 0.10) are highlighted in bold and italics.

| Parameters | Soil mycobiota composition |  |  | Wheat root endospheric mycobiota composition |  |
| --- | --- | --- | --- | --- | --- |
|  | df | F | P | F | P |
| Floristic richness | 1 | 0.68 | 0.961 | 1.49 | **0.021** |
| Residuals | 58 |  |  |  |  |
| Floristic evenness | 1 | 0.089 | 0.657 | 1.37 | **0.080** |
| Residuals | 58 |  |  |  |  |

### 3.2 Effects of weeds on wheat root endospheric mycobiota structure in the controlled experiment

Weed species were associated with distinct mycobiota composition from that found in wheat plants (Figure 1A). Mycobiota composition differed in the roots of each weed species (Figure 1B). Along with the first principal component of weed root mycobiota, the biggest differences were found between *P. rhoeas, T. repens;* and *V. sativa* (Figure S3A), while along with the second principal component of weed root mycobiota, the biggest difference was found between *M. chamomilla* and *G. aparine* (Figure S3B). *P. annua* and *P. trivialis* had the most similar root endospheric mycobiota composition along both principal components (Figure S3A-B). The effect of weed species was also significant when considering wheat root endospheric mycobiota, which clustered depending on the neighbourhood weed species they grew with (Figure 1C). Along with the first principal component of wheat root endospheric mycobiota, wheat root endospheric mycobiota differed the most between *P. rhoeas, M. chamomilla;* and *P. annua* treatments (Figure S3C), while along with the second principal component of wheat root endospheric mycobiota, *P. rhoeas* and *V. sativa* showed the biggest different effects (Figure S3D). But *P. annua* and *P. trivialis* did not have the same effect on wheat root endospheric mycobiota (Figure S3C-D).

**Figure 1.**
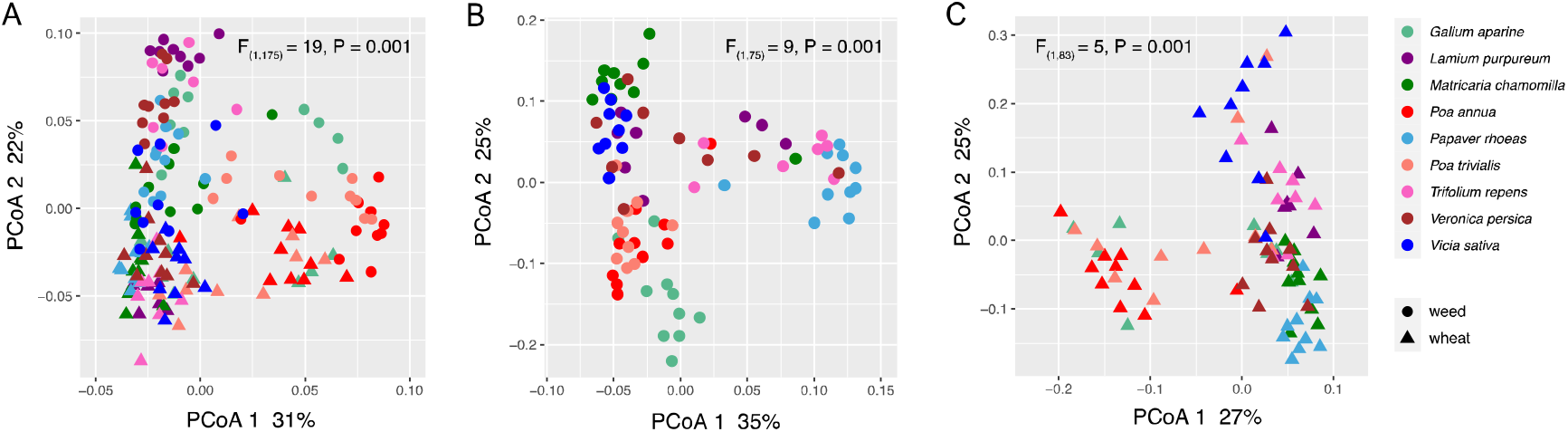
Composition of root endospheric mycobiota of weed species and wheat grown using plant-matrix design in the controlled experiment. (A) PCoA of root endospheric mycobiota of all weed and wheat plants; (B) PCoA of root endospheric mycobiota of all weed plants; (C) PCoA of root endospheric mycobiota of all wheat plants with different weed species as neighbours.

### 3.3 Effects of weeds on wheat root endospheric mycobiota diversity in the controlled experiment

The weed species *P. trivialis* displayed the highest root endospheric mycobiota sequence cluster richness, the weed species *V. persica* also displayed relatively high root endospheric mycobiota richness, while the two weed species *M. chamomilla* and *V. sativa* had the lowest sequence cluster richness (Figure 2A). Neighbourhood weed identity had a significant effect on the wheat root endospheric mycobiota sequence cluster richness (Table S3), in which the wheat individuals growing with *G. aparine, M. chamomilla, P. rhoeas, P. annua, P. trivialis, T. repens* and *V. persica* displayed significantly higher root mycobiota sequence cluster richness than individual wheat plants growing with *L. purpureum* (Figure 2B). The weed species *V. persica* displayed the highest root mycobiota evenness, while the weed species *P. annua* had the lowest root mycobiota evenness (Figure 2C). No significant differences were found in root endospheric mycobiota evenness among individual wheat plants growing with different weed species (Figure 2D).

**Figure 2.**
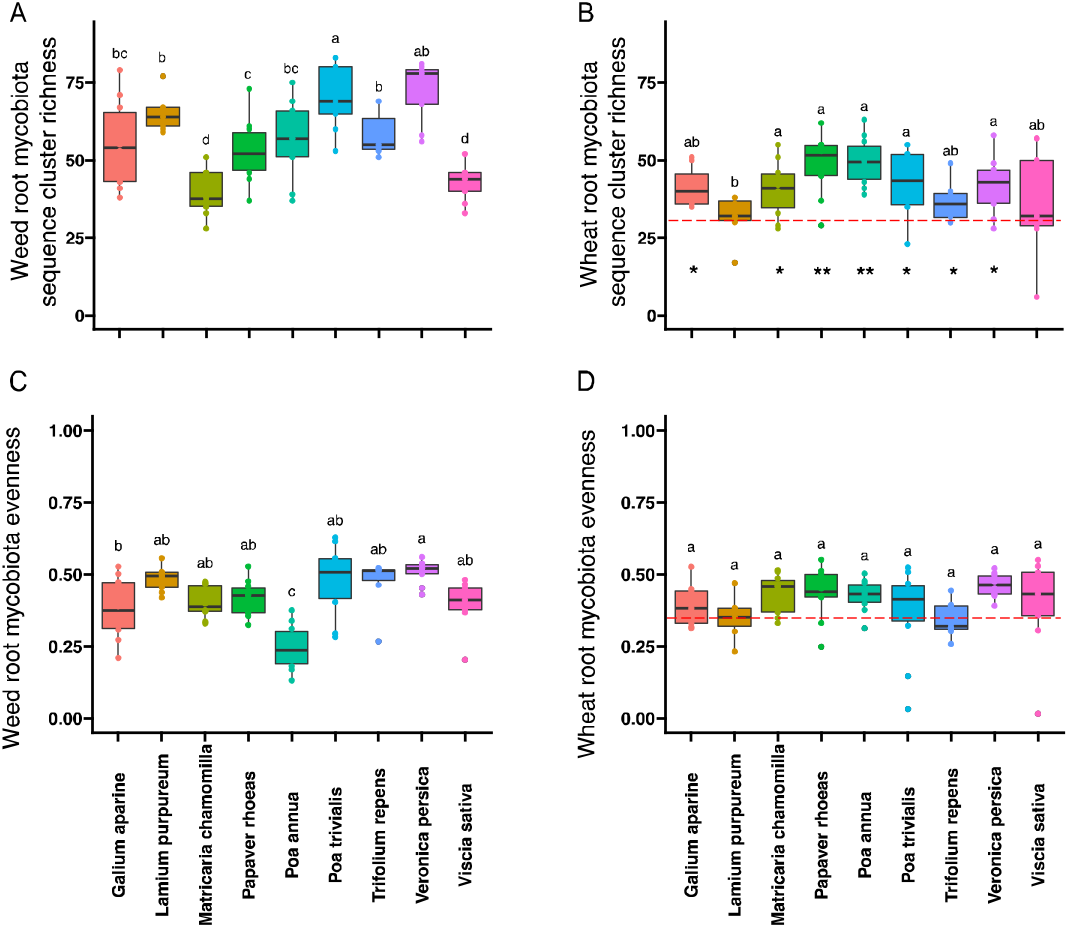
Weed and wheat root endospheric mycobiota sequence cluster richness in the controlled experiment. (A) Root endospheric mycobiota sequence cluster richness of the neighbouring weed species; (B) Root endospheric mycobiota sequence cluster richness of wheat plants. (C) Root endospheric mycobiota evenness of the neighbouring weed species; (D) Root endospheric mycobiota evenness of wheat plants. In (B) and (D), the red dashed line indicates, respectively, the mean root mycobiota sequence cluster richness and evenness of wheat individuals in the control treatment of a single wheat plant growing in the pot. Asterisks indicate the significance level of weeds in promoting wheat root mycobiota diversity compared with red dashed line: * indicates 0.01< P<0.05; ** indicates P < 0.01. Lowercase letters indicate significant differences in weed identity (Tukey post-hoc test) in all treatments.

### 3.4 Effects of weeds on their ability to transmit root endospheric mycobiota to wheat in the controlled experiment

Different weed species shared 10% to 70% sequence clusters (i.e. 5 to 45 sequence clusters) with wheat roots (Figure 3). *G. aparine*, *P. rhoeas*, *P. annua*, *P. trivialis, V. persica* and *V. sativa* shared the highest number of sequence clusters with wheat (Figure 3A), while *M. chamomilla* and *P. annua* shared the highest percentage of their own root endospheric mycobiota with wheat roots (Figure 3B). The smallest number and the lowest percentage of shared weed root endospheric mycobiota to wheat roots were found for *T. repens* and *L. purpureum*, respectively (Figure 3B).

**Figure 3.**
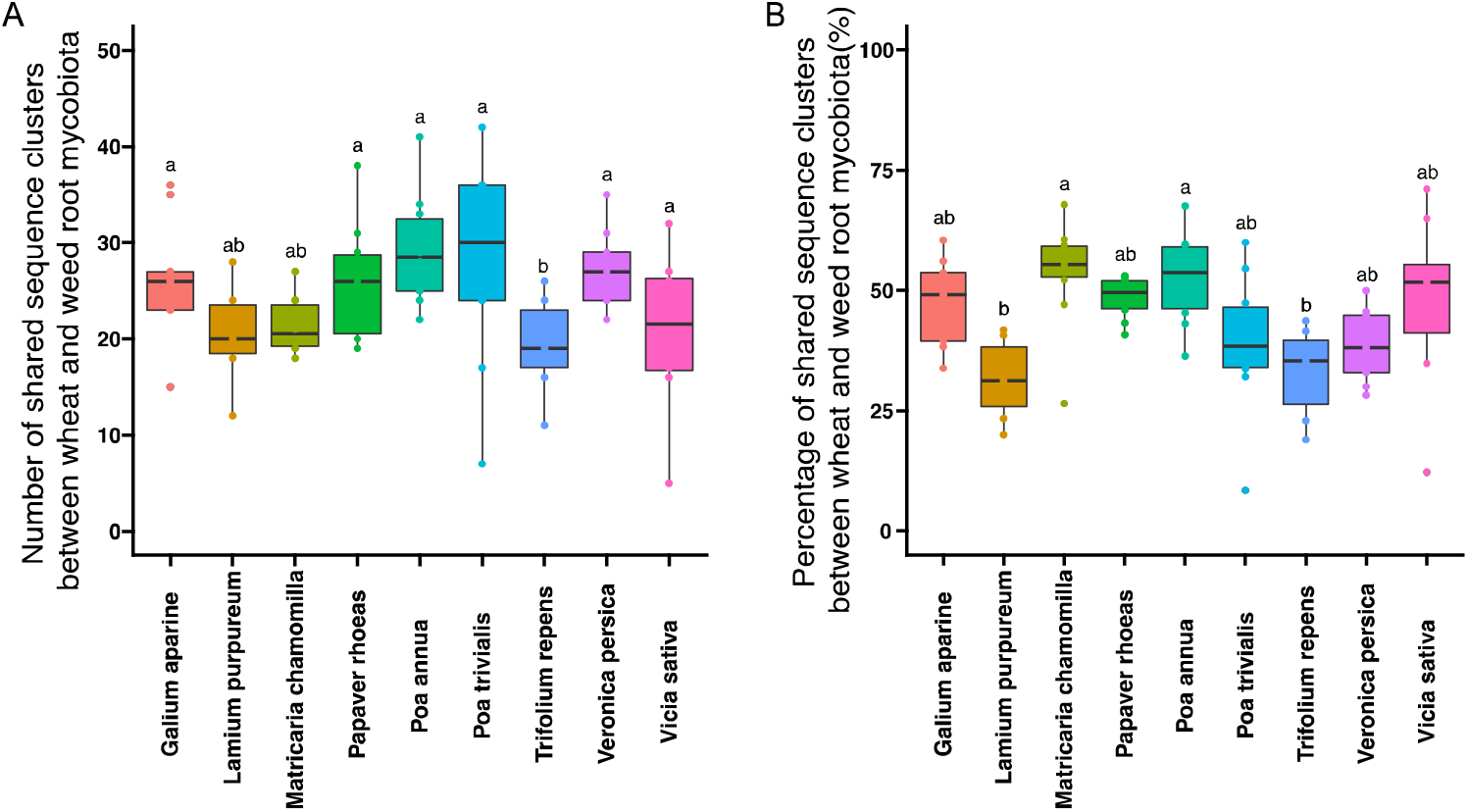
Root endospheric mycobiota transmission from neighbouring weed plants to focal wheat plants in the controlled experiment. (A) Number of shared root endospheric mycobiota sequence clusters between wheat and neighbouring weed plants; (B) Percentage of shared root endospheric mycobiota sequence clusters between wheat and neighbouring weed plants. Lowercase letters indicate significant differences in weed identity (Tukey post-hoc test) in all treatments.

In almost all cases, weed neighbourhoods enriched mycobiota in wheat microbiota compared to wheat alone, only a few sequence clusters were decreased. This enrichment was dependent on the neighbouring weed species (Figure 4A). *L. purpureum* and *P. annua* positively modified the relative abundance of the amount of 35 and 34 sequence clusters, respectively, while *T. repens* had the least influence on the wheat root endospheric mycobiota (Figure 4A). *P. trivialis* and *V. sativa* reduced the relative abundance of five sequence clusters, and this was the strongest negative effect on the root endospheric mycobiota of individual wheat plants (Figure 4A). Some sequence clusters (e.g. clusters 25 and 30, phylum Ascomycota) were transmitted successfully to wheat roots by most of the weed species, while other specific sequence clusters (e.g. clusters 16 and 432, species Geranomyces) were only transmitted successfully by one weed species (Figure 4B). This generalist versus specialist effect was more obvious in weed reduced clusters, cluster 23 (phylum Ascomycota, family Capnodiales) and cluster 5 (phylum Glomeromycota, genus Gigasporaceae) were reduced by most weed species, whereas cluster 85 (phylum Ascomycota, species Gloeotinia), cluster 10 (phylum Ascomycota, family Hypocreales) and cluster 16 (phylum Chytridiomycota, species Geranomyces) were only reduced by *Vicia sativa* (Figure 4C). Clusters belonging to Glomeromycota were reduced by weed species *L. purpureum, P. annua, P. rhoeas, P. trivialis and T. repens*, the relative abundance of most sequence clusters in the Ascomycota of wheat root mycobiota were increased by the presence of weeds (Figure S4).

**Figure 4.**
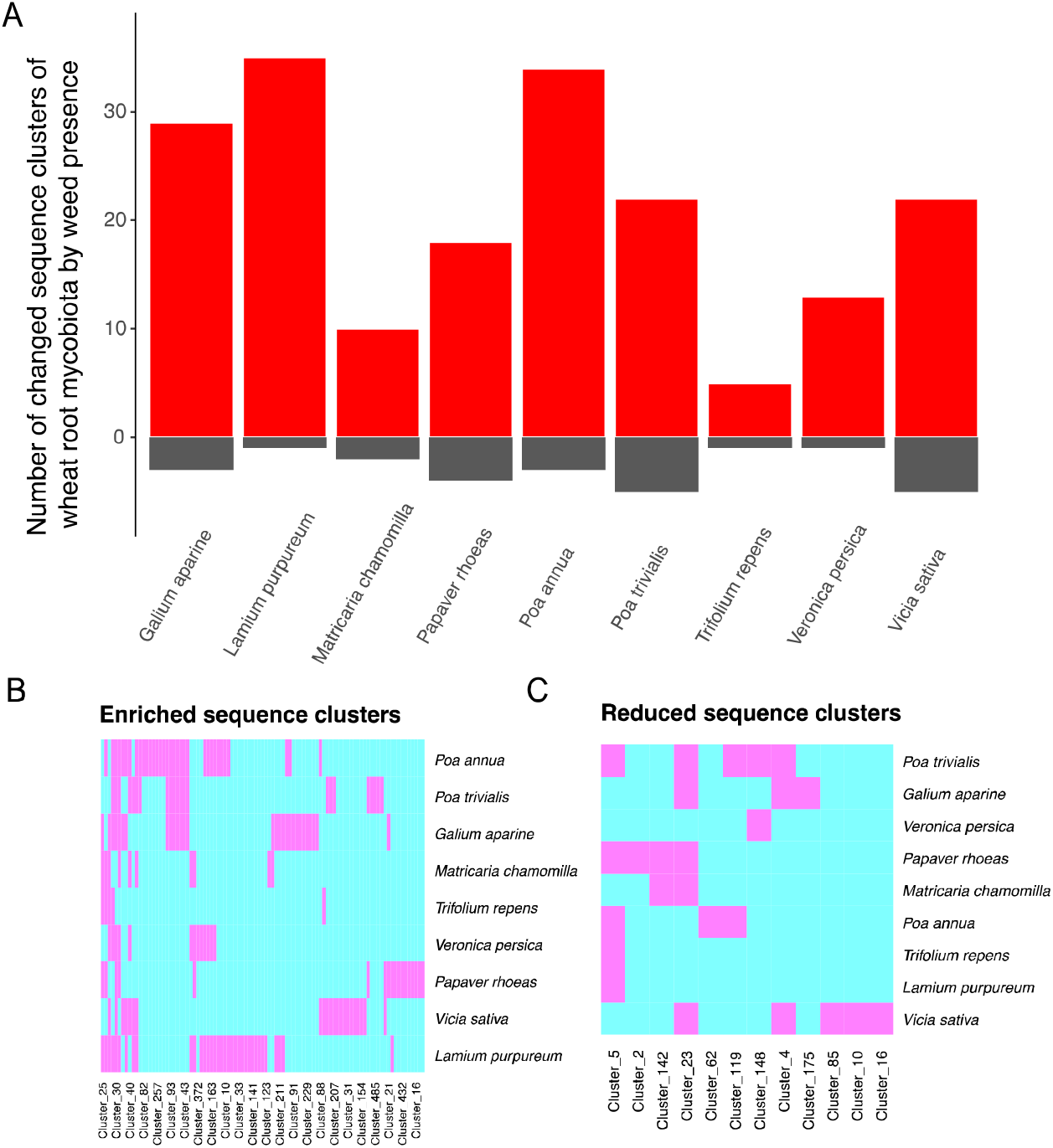
Effect of the neighbouring weed species on the relative abundance of sequence clusters associated with wheat in the controlled experiment. In the 3 panels, the neighbourhood effects are shown relative to the wheat only control. (A) Number of significantly (P < 0.05) modified sequence clusters in wheat root endosphere, the red bars indicate the enriched amount (i.e. relative abundance with log2FoldChange > 0.6) of root endospheric mycobiota sequence clusters and the grey bars indicate reduced amount (i.e. relative abundance with log2FoldChange < - 0.6) of root mycobiota sequence clusters; (B) Identity of enriched sequence clusters in wheat root endospheric mycobiota; (C) Identity of reduced sequence clusters in wheat root mycobiota. In both (B) and (C) the pink grids indicate significantly changed sequence clusters in the wheat root, either increased or decreased relative abundances.

### 3.5 Effects of weeds on wheat performance via their root endospheric mycobiota in the controlled experiment

In the controlled experiment, the aboveground dry biomass of neighbouring weeds varied depending on the weed species (Figure 5A). In all treatments with weeds as neighbours, wheat aboveground biomass was greater than that of wheat individuals surrounded by four wheat plants. In five out of the nine treatments with weeds as neighbours, treated wheat individuals had more aboveground biomass than the individual wheat plants growing alone. Neighbourhood weed identity had a significant effect on wheat aboveground biomass (Figure 5B, Table S3). *V. persica* and *V. sativa* not only gained growth by themselves but also showed the most improvement in wheat biomass compared to controls (Figure 5), whereas *P. rhoeas* gained in self growth (Figure 5A) but did not promote wheat growth (Figure 5B). The total number of sequence clusters, especially those related to Ascomycota and Basidiomycota, associated with wheat root endospheric mycobiota significantly increased wheat aboveground biomass (Table 3).

**Figure 5.**
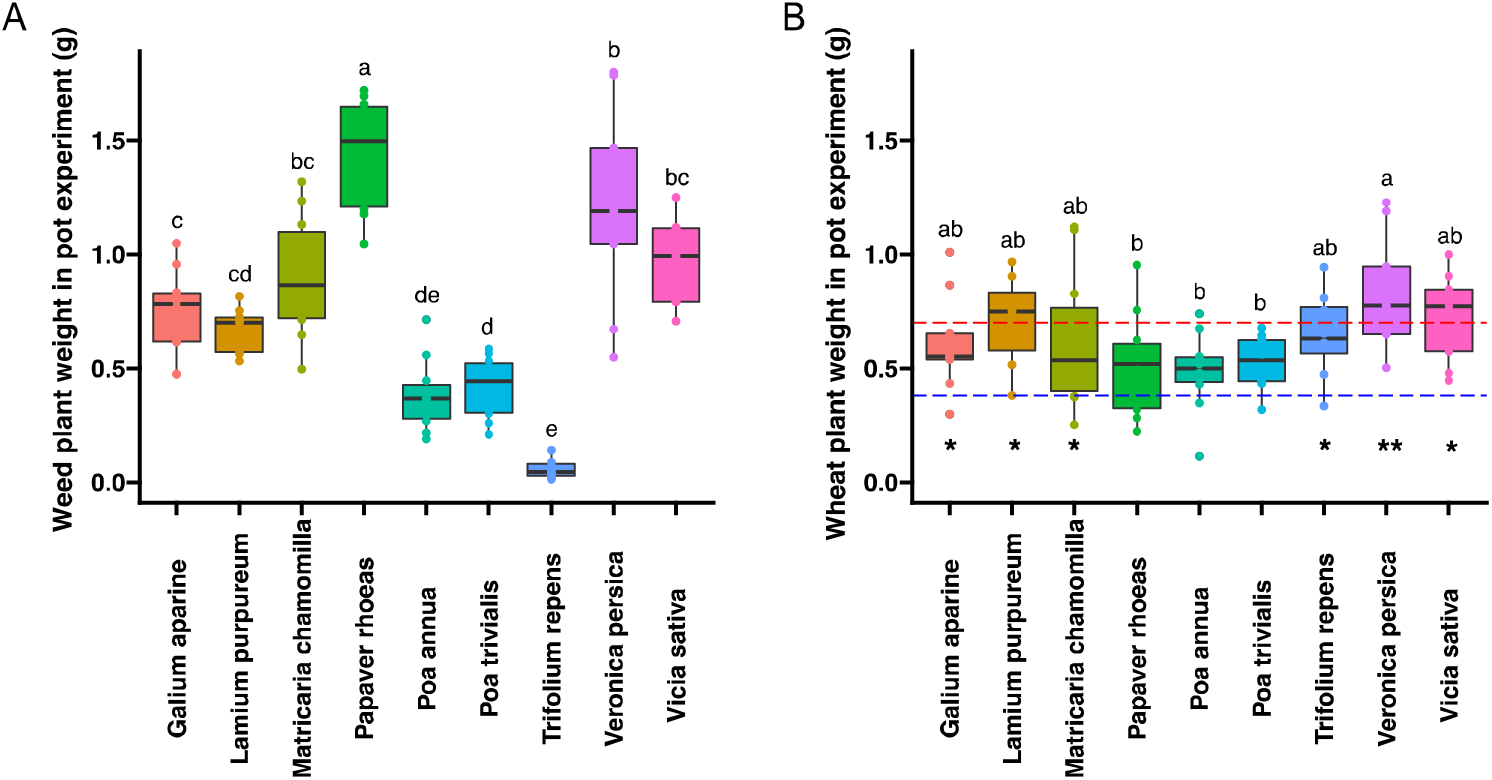
Plant aboveground biomass of different weed species and wheat in the controlled experiment. (A) Aboveground biomass of weed species; (B) Wheat aboveground biomass depending on the neighbouring weed species. In (B), the red and the blue dashed lines indicate the mean biomass of wheat individuals in the control treatment in which a single wheat plant was grown in each pot and in the control treatment in which the wheat plant in each pot was surrounded by four individual wheat plants respectively. Asterisks indicate the significance level of weeds in promoting wheat growth compared with blue dashed line: * indicates 0.01< P<0.05; ** indicates P < 0.01. Lowercase letters indicate significant differences in weed identity (Tukey post-hoc test) in all treatments.

**Table 3.** Effect of different predictors on wheat performance (aboveground dry biomass) in the controlled experiment. The predictors included sequence cluster richness of whole wheat root mycobiota, Ascomycota, Basidiomycota, Chytridiomycota, Glomeromycota and Zygomycota. Significant results (P < 0.05) are highlighted in bold, and marginal significant results (0.05 < P < 0.10) are highlighted in bold and italics.

| Parameters | df | Chisq | P |
| --- | --- | --- | --- |
| Wheat root mycobiota richness | 1 | 3.59 | **0.06** |
| Ascomycota sequence cluster richness | 1 | 3.13 | **0.08** |
| Basidiomycota sequence cluster richness | 1 | 5.51 | **0.02** |
| Chytridiomycota sequence cluster richness | 1 | 1.44 | 0.23 |
| Glomeromycota sequence cluster richness | 1 | 1.29 | 0.26 |
| Zygomycota sequence cluster richness | 1 | 0.40 | 0.53 |
| Residuals | 73 |  |  |
| Model summary | $R^2 = 0.11$ | AIC = 27 | |

## 4 Discussion

### 4.1 Weed neighbours enriched and shaped composition of wheat microbiota but not by modifying soil mycobiota

We showed that the composition and richness of neighbouring weeds influenced the mycobiota associated with wheat roots (Table 2), whereas little effect was found on bulk soil mycobiota. This suggests that the observed neighbourhood effects of weed plants on the mycobiota of wheat plants were likely due to local microbial dispersal from neighbour plants to crop plants rather than to a change in soil microbial reservoir in which the crop plant recruits. This result contrasts with that obtained in a previous study showing that a neighbour effect led to a legacy effect (Vannier et al., 2020) when the plant communities had been growing in the soil for several years. In the present study, the limited legacy effect was probably due to the short lifespan of the weeds, which were annual plants and only grew in soil for a maximum of one year, as the soil was plowed in preparation for transplantion the wheat plantlets. Local transmission of fungi among plants has already been demonstrated in an experiment performed to test the effect of plant neighbours on *Medicago truncatula* (Mony et al., 2021). Processes of microbial transmission between plants can be achieved by microbial inoculation via contact between roots or leaves (Enkhtuya et al., 2005; Smith & Read, 2008) or by the development of hyphae (Simard, 2018).

In addition, we demonstrated a positive effect of the diversity of weed neighbours on wheat root endospheric mycobiota diversity in most fungal phyla including Ascomycota, Glomeromycota and Zygomycota (Table S2). Because plants are associated with a preferential mycobiota (*sensu* host-preference effect (Vandenkoornhuyse et al., 2002)), diverse plant communities provide a higher diversity of niches for microorganisms, thereby encouraging a bigger range of microorganisms to coexist locally. Some evidence has shown that richer plant communities increased the diversity of total plant microbiota associated with the shoot (Navrátilová et al., 2018). The increased diversity of fungi provided by a diverse neighbourhood is a possible reservoir for transmission to crop plants growing nearby. Here, we demonstrated that, despite an existing soil microbiota that harboured much higher diversity than the microbiota associated with plants, weed neighbourhoods, even with less abundant cover, significantly influence the root-associated mycobiota of crop plants growing close by (i.e. at a distance of less than one meter).

### 4.2 Weed neighbours affected wheat root endospheric mycobiota and wheat performance

We assessed the ability of neighbourhood plants to influence the wheat root endospheric mycobiota in controlled conditions. Using nine different weed species cultivated in organic field soil as inoculum for wheat plants growing in sterile conditions, we observed differences in the ability of weed species to recruit their own root endospheric mycobiota, and to manipulate the wheat root microbiota. In weeds, these processes, which were linked to the difference in the influence of a target neighbouring plant can be explained in three steps. Firstly, weed species differ in their ability and use different patterns to recruit root mycobiota, as already shown for arbuscular mycorrhizal fungi (AMF) by (Vatovec et al., 2005), who classified 14 weed species as strong, weak and non-host plants for AMF. Plant phylogeny plays a role in structuring their root microbiomes, and a previous study showed that plants that are distantly related phylogenetically show greater variation in the composition of their associated microbiome (Bouffaud et al., 2014). In the present study, the composition of root endospheric mycobiota of phylogenetically similar weed species such as *P. annua* and *P. trivialis, T. repens* and *V. persica*, was also similar. Secondly, plant root exudate profiles may also influence the recruitment of root endospheric mycobiota (Pascale et al., 2020; Voges et al., 2019) and their surrounding microorganisms thereby creating a unique microbial reservoir for their neighbouring plants. Thirdly, plant root traits could explain the transmission of root mycobiota to plant neighbours, as it has been **Error! Bookmark not defined.**shown that neighbourhood plants’ functional proximity in terms of belowground resource use and uptake strategy was a key predictor of a neighbouring effect on focal plants (Mony et al., 2021).

The ability of weed species to transmit their root endospheric mycobiota to nearby wheat roots also depends on the species. We demonstrated that some sequence clusters were transmitted to wheat roots by most of the weed species tested, including clusters belonging to the Ascomycota phylum. Conversely, some sequence clusters were specifically transmitted by particular weed species to wheat roots. For example one species of *Geranomyces* belonging to the Chytridiomycota phylum described as parasites of arbuscular mycorrhizae (Simmons, 2011; Wakefield et al., 2010), were only transmitted by *P. rhoeas*. By manipulating this Geronomyces species, *P. Rhoeas* might improve its own competitive advantage. Future studies on the functions of this neighbour driven microbiota manipulation are required.

As expected, we demonstrated competitive interactions between wheat individuals and neighbouring weeds. However, some weed species promoted wheat growth compared to the ‘wheat grown alone’ control (e.g. *Veronica persica*). In our experimental design, the wheat growth promotion was necessarily mediated by neighbouring weed species and this phenomenon was shown to be correlated with modifications in wheat mycobiota. Among the weed species studied here, some were particularly beneficial for wheat growth (e.g. *Veronica persica*, *Vicia sativa* or *Matricaria sp.*) relative to their competitive influence. Modifications to wheat mycobiota caused by weed neighbours could increase crop yield if their effects on wheat biomass are confirmed in field conditions.

### 4.3 Weeds as auxiliaries for crops in sustainable agricultural system

Intensive agriculture has led to a major reduction in soil diversity (Tsiafouli et al., 2015), and disrupted the plant-microbial symbiosis (Porter & Sachs, 2020). In particular, the long history of plant breeding has reduced the ability of domesticated crop plants to efficiently recruit their own microbiome from surrounding microbial reservoirs. Knowing that wheat breeding may have resulted in wheat plants that are no longer able to efficiently filter or recruit their microbiota endosphere (Mauger et al., 2021), wild neighbour auxiliary plants might be able to mitigate the disturbance of the wheat microbiota in modern crops through their influence on wheat microbiota.

The role of weed plants in agrosystem functioning is already known (Gaba et al., 2020; Marshall et al., 2003), for instance, affecting the composition and interactions of the insect fauna to protect beneficial insects, thereby increasing pollination, providing microclimates for crop development, and regulating the development of competitive weeds. Beyond these ecological functions, in the present study, we demonstrated that weed species can help enrich plant microbiota and transmit specific sequence clusters when they grow in the close neighbourhood of crop plants. Such neighbourhood effects, which are likely caused by root-root connections, can also transmit systemic acquired resistance against pathogens to neighbouring plants (Cheol Song et al., 2016), thus helping plants survive and adapt to different environments. Weed neighbourhood effects require in-depth analysis in both controlled and field conditions including screening larger sets of neighbouring weed species in order to identify candidate auxiliary weed plant species that could be promoted in crops through dedicated field management.

Agricultural management is not ‘all rocket science’ (“Agriculture Isn’t All Rocket Science,” 2021). Contrary to the current direction, developing ecological approaches to agriculture can be used to favour future sustainable food production, all elements in agricultural systems including both plant diversity, their associated microbiota and the soil microbial reservoir shall be taken into consideration for a more holistic agriculture to obtain higher and more stable crop yields in a more sustainable way. In this context, weed plants could be used as auxiliary plants that provide ecosystem services to targeted crop plants in agricultural systems.

## Author Contributions

C.M. and P.V. conceived the study and methodology. C.R., J.H. and P.F. collected the data and performed the sequence analyses for weed and wheat root mycobiota, S.M. processed and performed the sequence analyses for soil samples. J.H. and C.R. performed statistical analysis. J.H. and C.R. wrote the manuscript with the help of C.M. and P.V. All the authors contributed to the manuscript and gave approval for publication.

## Data Availability Statement

Sequence data are deposited in the Sequence Read Archive (SRA) under accession number PRJNA811118. Data and scripts relevant to this manuscript are available at https://github.com/HuJamie/Weed-neighbourhood-effect-on-wheats.

## Acknowledgements

We thank the GenoSol platform and the EcogenO platform, both are part of ANAEE-France research infrastructure and Biogenouest. GenoSol received a grant from the French National Agency for Research (ANR-11-INBS-0001) and a grant from the Regional Council of Bourgogne-Franche-Comté. The BRC GenoSol is a part of BRC4Env, the pillar “Environmental Resources” of the Research Infrastructure AgroBRC-RARe. J.H. was supported by a grant from the Brittany council for H2020-MSCA-IF projects 2019 Seal of Excellence - CORRiBIOM, and C.R. was supported by a grant from Fondation de France, *l’Agence Française pour la Biodiversité* and a grant from PIA ANR DEEP IMPACT (ANR-19-CPA-00XX-05).

