## supplemental materials for "Neighbourhood effect of weeds on wheat root endospheric mycobiota"

**Supplementary materials and methods**

**Experimental design**

In the field study, after sampling in the fields, soil and root samples were carefully stored and processed in the following 24 hours, after which the soil and wheat root endospheric microbiota were analysed by metabarcoding.

In the controlled experiment. First, each weed species was pre-cultivated in an organic field soil for four weeks. Weed seeds were germinated and grown in separate pots filled with an organic field soil (pH: 6.8; MO: 3.3%; P_2_O_5_: 85 mg/kg; C/N: 8.4). The soil was sampled from one organic field in the eastern part of the LTSER area belonging to the set that was analysed in the first study. Soil was sieved through a 5-mm mesh and homogenised before the controlled experiment. Pots containing weed plantlets were placed in a growth chamber (21 ± 0.5 °C; 12L:12D; 10000 lux) and watered with sterile water every two days and with a sterile 0.5X Hoagland solution every two weeks. After four weeks, weed plantlets were collected and washed with sterile water to remove soil particles and rhizospheric microorganisms attached to the roots. In parallel, individual wheat plants were cultivated for two weeks until the second node stage (21 ± 0.5°C; 12L:12D; 5200 lux) then vernalised in sterile conditions for two months (4 ± 0.5 °C; 8L:16D; 10000 lux). One sterile individual wheat plant surrounded by four individuals of the same weed species was planted in 3 L pots (diameter 19 cm, height 15 cm) filled with sterile vermiculite substrate. The sterile wheat plant was pre-cultured in organic soil containing endospheric mycobiota (Figure S1). Neighbouring plants were planted at similar distances to the focal wheat plant. The pots were randomly placed in a controlled growth chamber (21 ± 0.5 °C; 12L:12D; 5200 lux). The pots were watered with a standardised quantity of sterile demineralised water (100 mL, 150 mL and 200 mL per week depending on the progress of the experiment) and 100 mL of sterile 0.5X Hoagland solution. All the plants were harvested 13 weeks after transplantation.

**DNA extraction**

Briefly, 250 mg of lyophilized soil were used for DNA extraction (QIAGEN DNeasy PowerSoil kit). The quality and the integrity of the soil DNA were checked by electrophoresis on 1% agarose gel. DNA was quantified by measuring PicoGreen fluorescence using a Quant-iT™ dsDNA Assay Kit (Invitrogen, USA). Briefly, 80 mg of root samples were ground to a powder, DNA was extracted using magnetic beads (sbeadex mini plant kit, LGC genomics) following the manufacturer's protocol and using a robot (oKtopure robot, LGC Genomics). DNA quality was checked by running samples on 1% sodium boric acid agarose gel electrophoresis, and DNA concentrations were measured by fluorimetric quantification (Hoeschst). The concentrations of all the qualified DNA samples were normalized at 7 ng/μL (Bravo-Agilent®) and 21 ng of DNA was used for each PCR. Extracted DNA was stored at -80 °C for subsequent analyses.

**18S rRNA amplicon sequencing and bioinformatics**

Primers NS22b (5′-AATTAAGCAGACAAATCACT-3′) and SSU817 (5′-TTAGCATGGAATAATRRAATAGGA-3′) were used for specific amplification of the fungal V4 and V5 18S rRNA gene region (Borneman & Hartin, 2000; Lê Van et al., 2017) leading to a mean 530 pb amplicon. For multiplexing at their 3’ region, all the primers contained the Illumina® adaptors. The PCRs were performed with Illustra PuReTaq Ready-to-go beads (GE Healthcare®). The PCR programme comprised an initial denaturation step at 95 °C for 4 mins followed by 35 cycles at 95 °C for 30 s, 53.5 °C for 30 s, 72 °C for 1 min and a final extension step at 72 °C for 10 mins. A second PCR was performed using the Smartchip-system (Takara) to achieve multiplex tagging (up to 384 amplicons in a one-step PCR). The tagged-amplicon pool was then purified (AMpureXP, Agencourt®) and quantified using Kapa Library Quantification Kit- Illumina® platforms (KAPA BIOSYSTEMS®) on a LC480 LightCycler qPCR instrument (Roche®). Pair-End 2x300 cycles sequencing runs (MiSeq instrument, Illumina) and sequencing libraries were performed. The DNA concentration of each sample was normalized, the amplicon library constructed and finally, sequencing was performed.

SWARM is an adaptive sequence agglomeration based on aggregation parameters rather than on a global similarity threshold (Mahé et al., 2014) to produce “sequence clusters”. Compared to the ASV approach (Callahan et al., 2016), the main advantage of clustering in FROGS is avoiding over-estimation of sequence cluster diversity (e.g. multicopy artifacts). Sequence clustering was combined with a rigorous chimera removal step (Escudié et al., 2018) followed by a step to remove all sequence clusters detected in less than three independent samples and with a threshold of 0.005% of reads in all. Sequence clusters were filtered using the quality of the affiliations with a threshold of at least 95% BLAST identity and 95% coverage.

**Supplementary results**

*Juncus bufonius* (50 neighbourhoods), *Trifolium repens* (49 neighbourhoods), *Poa annua* (48 neighbourhoods) and *Matricaria sp.* (45 neighbourhoods) were present in most surveyed neighbourhoods (Table S1). 27 out of the 96 weed plant species were present in fewer than three out of 60 neighbourhoods and were not used for subsequent analysis. The soil mycobiota and wheat root endospheric mycobiota was composed of 467 and 423 sequence clusters, respectively, dominated by Ascomycota, followed by Zygomycota and Basidiomycota, lastly by Chytridiomycota and Glomeromycota (Figure S5). Soil mycobiota sequence cluster richness and evenness were both significantly higher than those of wheat root endospheric mycobiota (Figure S6). The structure of the field soil mycobiota was distinct from that of the wheat root endospheric mycobiota with greater distances between samples detected in root endospheric mycobiota composition than in soil with a denser cluster (Figure S7A). Soil shared 421 mycobiota sequence clusters with wheat root, and soil had 43 unique mycobiota sequence clusters, while wheat root only had three unique mycobiota sequence clusters (Figure S7B).

The wheat and weed root endospheric mycobiota were composed of 283 sequence clusters, including 56% of Ascomycota (101 sequence clusters), 3% of Basidiomycota (40 sequence clusters), 30% of Chytridiomycota (95 sequence clusters), 11% of Glomeromycota (20 sequence clusters) and 1% of Zygomycota (18 sequence clusters) (Figure S8).

**Supplementary figures**

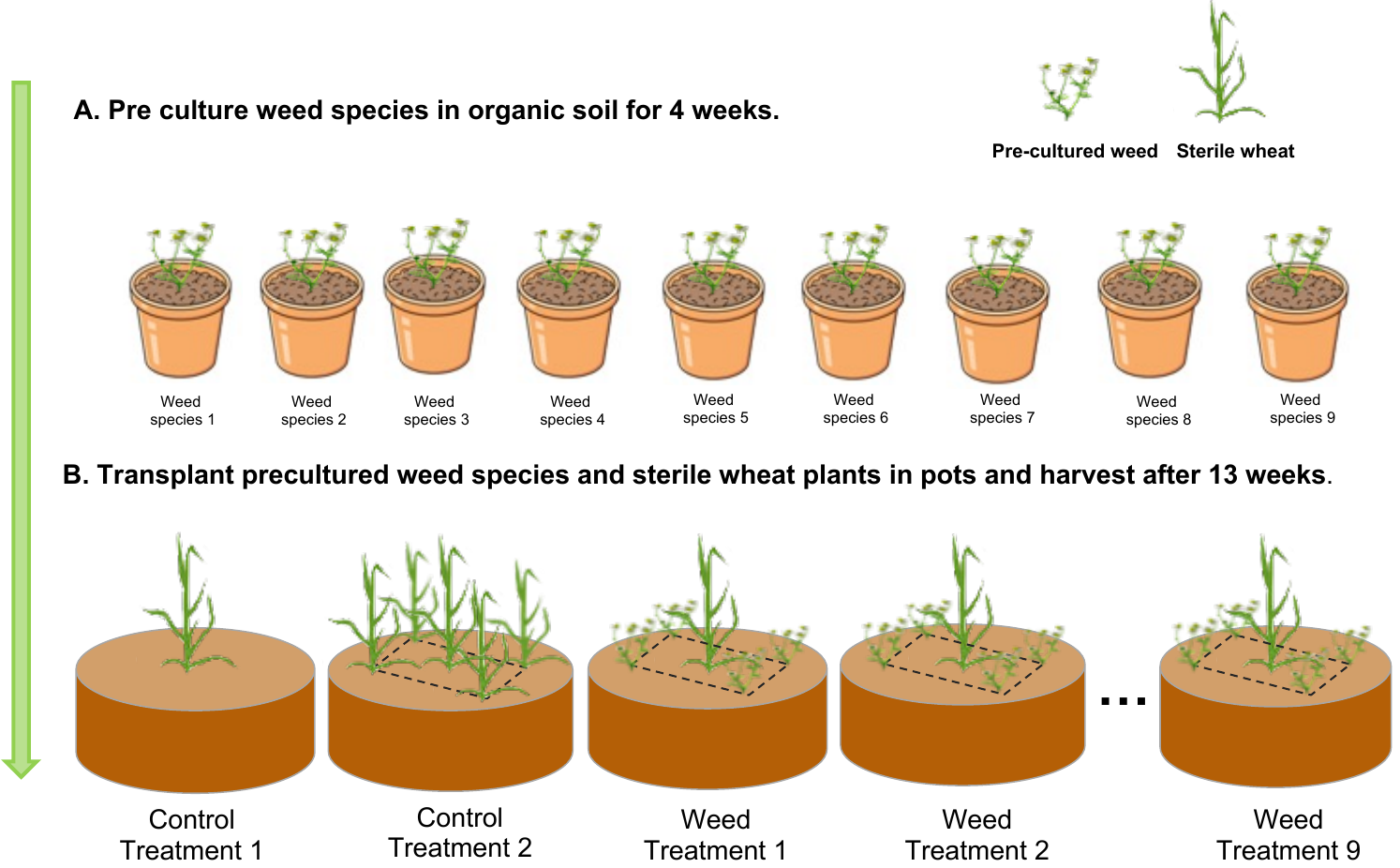

**Figure S1. Experimental design.** (A) pre-culture of weed species in soil sampled in an organic field; (B) Plant-matrix design, a sterile pre-vernalised wheat plant was transplanted to the centre of the pot, and four pre-cultivated individual weed plants (of the same species) were transplanted around it at the same distance from the focal wheat plant. Control treatment 1 contained one individual wheat plant growing alone in one pot. Control treatment 2 contained one individual wheat plant surrounded by four neighbouring wheat plants.

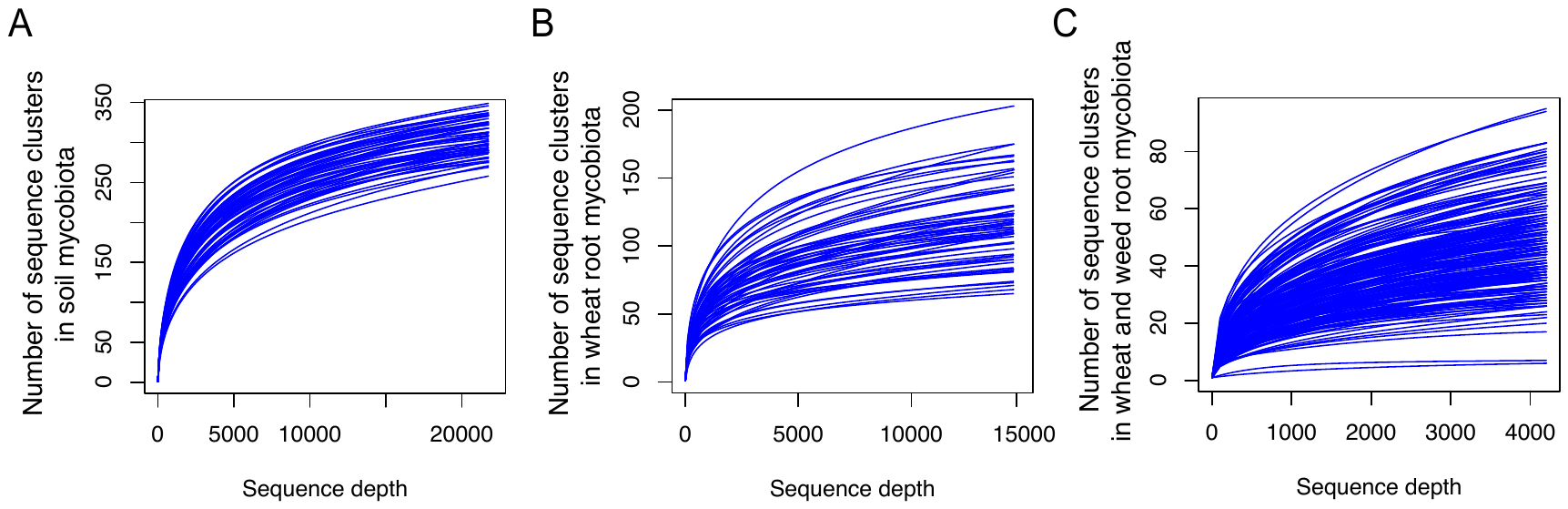

**Figure S2. Rarefaction curves of soil, wheat and weed root endospheric mycobiota.** Curves show the total assigned sequence clusters detected relative to the number of sequences in the soil sampled from field (A), in wheat roots sampled from field (B) and in the roots of wheat and weeds in the controlled experiment (C).

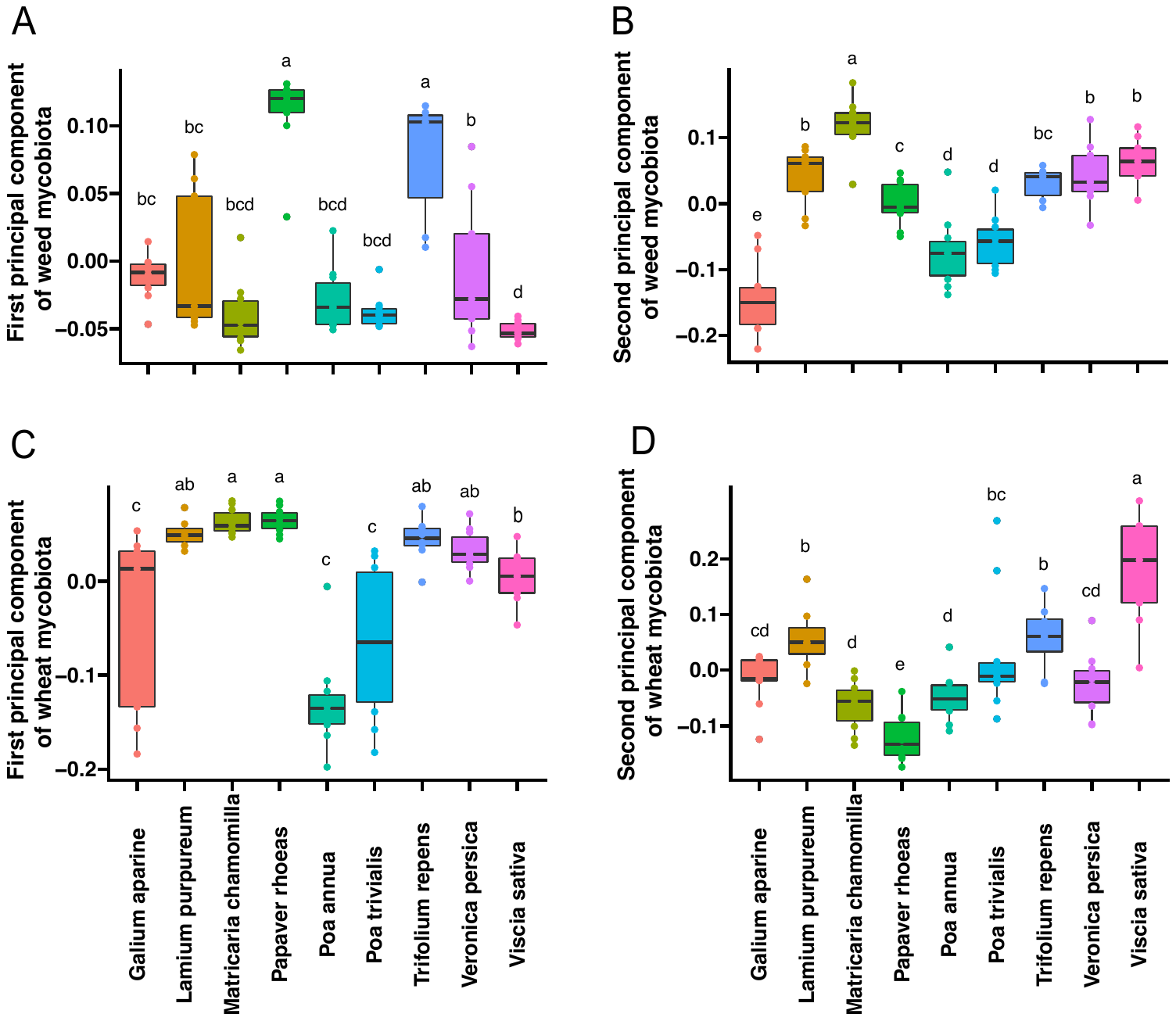

**Figure S3. Effect of different weed species on the first and second principal components of weed and wheat root endospheric mycobiota.** (A) Effect of weed species on the first principal component of the weed mycobiota; (B) Effect of weed species on the second principal component of the weed mycobiota; (C) Effect of weed species on the first principal component of the wheat mycobiota; (D) Effect of weed species on the second principal component of the wheat mycobiota.

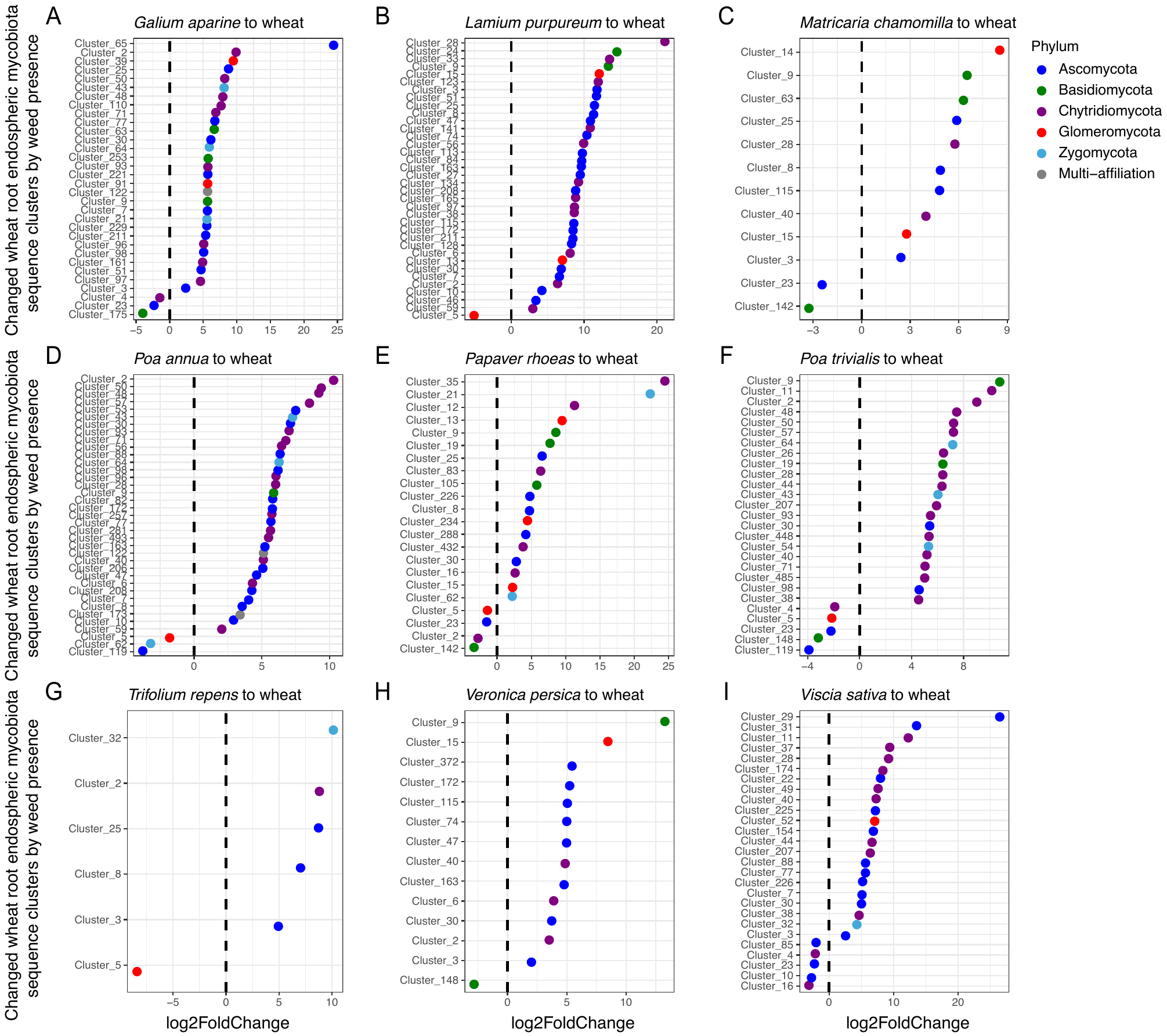

**Figure S4. Changes in the relative abundance of sequence clusters in wheat root mycobiota depending on the weed species in the controlled experiment.** The dashed black lines in all panels correspond to the threshold value of no change. The sequence clusters on the left side of the vertical dashed line were reduced and the sequence clusters on the right side of the vertical dashed line were enriched by the influence of the neighbouring weeds considered. The colour of different dots shows the taxonomic identification of the sequence clusters.

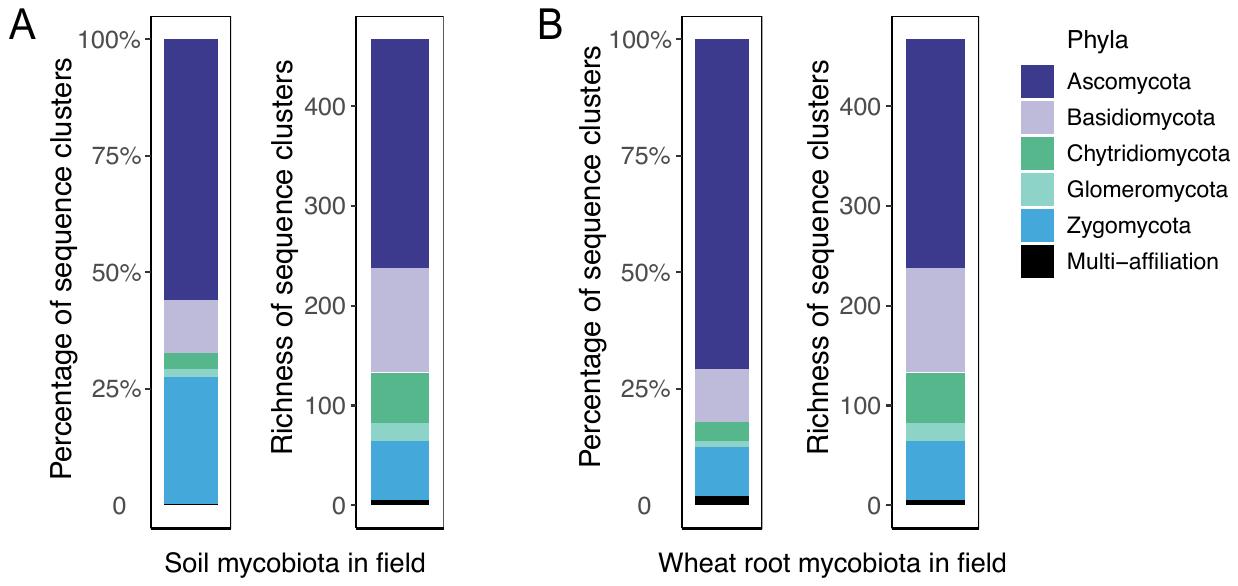

**Figure S5. Soil mycobiota and wheat root endospheric mycobiota sequence clusters in the field study.** (A) left panel: proportion of sequence cluster abundance in each phylum (Ascomycota, Basidiomycota, Chytridiomycota, Glomeromycota and Zygomycota) of soil mycobiota, (A) right panel: sequence cluster richness of each phylum (Ascomycota, Basidiomycota, Chytridiomycota, Glomeromycota and Zygomycota) in the soil mycobiota; (B) left panel: proportion of sequence cluster abundance in each phylum (Ascomycota, Basidiomycota, Chytridiomycota, Glomeromycota and Zygomycota) of wheat endospheric root mycobiota, (B) right panel: sequence cluster richness of each phylum (Ascomycota, Basidiomycota, Chytridiomycota, Glomeromycota and Zygomycota) in the wheat endospheric root mycobiota.

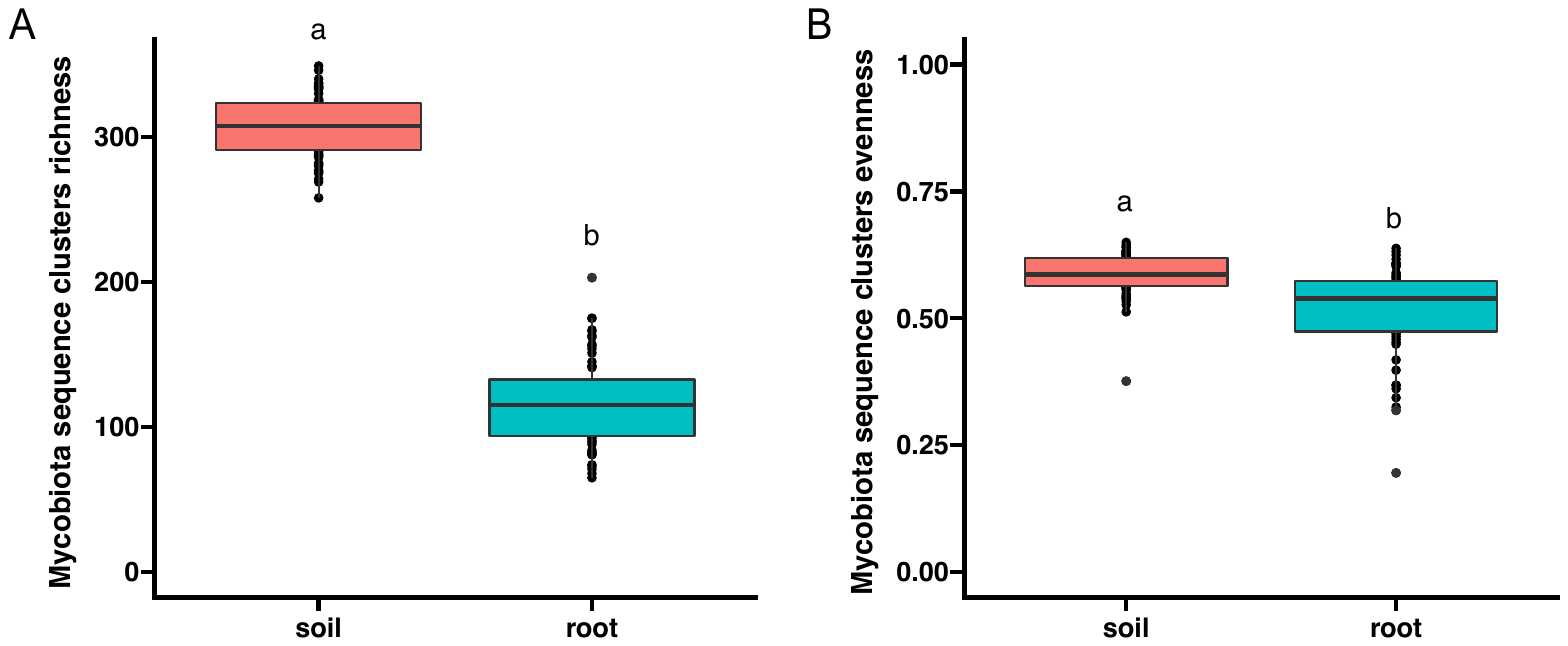

**Figure S6. Diversity of soil and wheat root endospheric mycobiota in the field study.** Mycobiota diversity is indicated as sequence cluster richness (A), and Pielou’s evenness index of the sequence cluster (B). Lowercase letters indicate significant differences between soil and wheat root mycobiota (t-test).

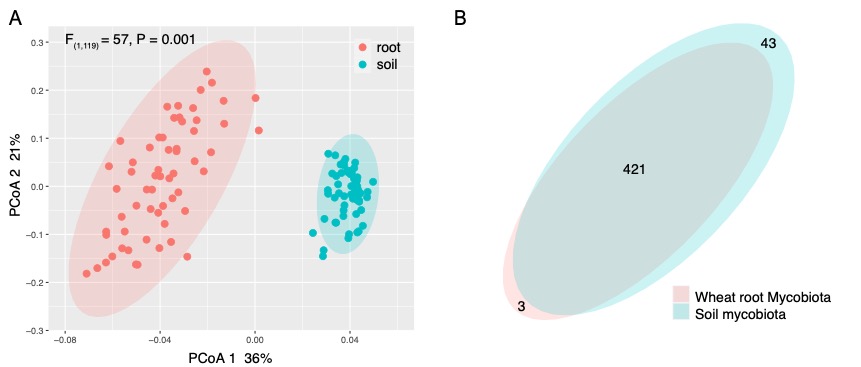

**Figure S7. Mycobiota composition and dissimilarity of soil and wheat root samples in the field study.** (A) PCoA of soil and wheat root mycobiota in the field study; (B) A Venn diagram of soil and wheat root mycobiota in the field study.

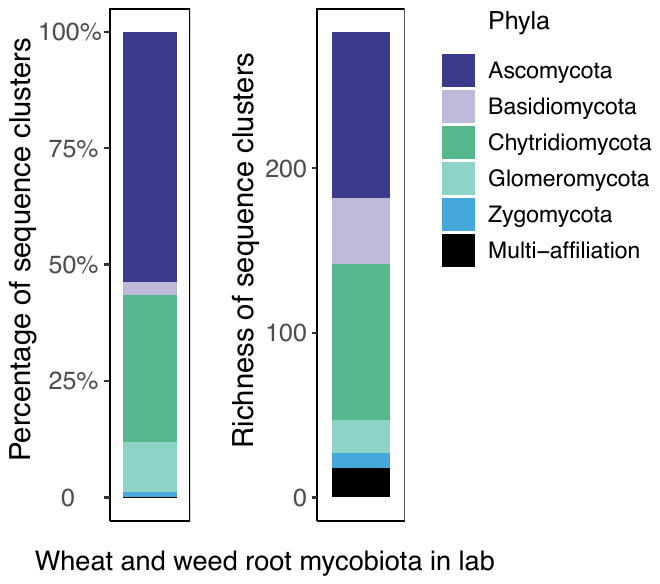

**Figure S8. Weed and wheat root endospheric mycobiota sequence clusters in the controlled experiment.** Left panel: proportion of sequence cluster abundance in each phylum (Ascomycota, Basidiomycota, Chytridiomycota, Glomeromycota and Zygomycota) in all weed and wheat root mycobiota; Right panel: sequence cluster richness in each phylum (Ascomycota, Basidiomycota, Chytridiomycota, Glomeromycota and Zygomycota) of all weed and wheat root mycobiota.

**Supplementary tables**

**Table S1. Occurrence, and mean coverage percentage of weed species in the neighbourhoods in the field study. Species used for the controlled experiment are highlighted in bold (n=60).**

| **Weed species** | **The number of present neighbourhoods** | **Mean of coverage percentage across all neighbourhoods** | **Mean of coverage percentage across presented neighbourhoods** |
| --- | --- | --- | --- |
| *Juncus bufonius* | 50 | 0.0880 | 0.1056 |
| ***Trifolium repens*** | 49 | 0.0567 | 0.0695 |
| ***Poa annua*** | 48 | 0.0656 | 0.0819 |
| ***Matricaria sp.*** | 45 | 0.0635 | 0.0847 |
| *Cerastium fontanum* | 42 | 0.0257 | 0.0367 |
| *Lysimachia arvensis* | 40 | 0.0582 | 0.0872 |
| *Veronica arvensis* | 37 | 0.0378 | 0.0612 |
| ***Poa trivialis*** | 36 | 0.0638 | 0.1063 |
| *Aphanes arvensis* | 31 | 0.0283 | 0.0548 |
| *Stellaria media* | 27 | 0.0252 | 0.0559 |
| ***Vicia sativa*** | 26 | 0.0647 | 0.1494 |
| *Taraxacum officinale* | 25 | 0.0259 | 0.0621 |
| *Capsella bursa-pastoris* | 24 | 0.0287 | 0.0719 |
| *Viola arvensis* | 23 | 0.0259 | 0.0676 |
| *Avena fatua* | 22 | 0.0634 | 0.1730 |
| *Convolvulus arvensis* | 22 | 0.0610 | 0.1664 |
| *Vicia hirsuta* | 21 | 0.0994 | 0.2839 |
| *Polygonum aviculare* | 20 | 0.0121 | 0.0363 |
| ***Veronica persica*** | 18 | 0.0161 | 0.0536 |
| *Chenopodium album* | 13 | 0.0076 | 0.0349 |
| *Epilobium tetragonum* | 13 | 0.0056 | 0.0257 |
| ***Papaver rhoeas*** | 13 | 0.0388 | 0.1789 |
| *Lolium multiflorum* | 12 | 0.0206 | 0.1031 |
| *Ranunculus repens* | 12 | 0.0204 | 0.1019 |
| *Arabidopsis thaliana* | 10 | 0.0034 | 0.0201 |
| *Arrhenatherum elatius* | 10 | 0.0349 | 0.2093 |
| *Avena sativa* | 10 | 0.0105 | 0.0628 |
| *Briza minor* | 10 | 0.0117 | 0.0704 |
| *Cirsium arvense* | 10 | 0.0613 | 0.3676 |
| *Trifolium pratense* | 10 | 0.0076 | 0.0458 |
| ***Galium aparine*** | 9 | 0.0077 | 0.0512 |
| *Oxalis fontana* | 9 | 0.0057 | 0.0379 |
| *Agrostis stolonifera* | 8 | 0.0164 | 0.1233 |
| *Cardamine hirsuta* | 8 | 0.0047 | 0.0353 |
| *Daucus carota* | 8 | 0.0072 | 0.0537 |
| *Geranium dissectum* | 8 | 0.0051 | 0.0386 |
| *Polygonum persicaria* | 8 | 0.0050 | 0.0378 |
| *Rumex obtusifolius* | 8 | 0.0028 | 0.0212 |
| *Centaurae nigra* | 6 | 0.0108 | 0.1083 |
| *Crepis capillaris* | 6 | 0.0021 | 0.0208 |
| *Crepis setosa* | 6 | 0.0068 | 0.0685 |
| *Kickxia elatine* | 6 | 0.0030 | 0.0298 |
| ***Lamium purpureum*** | 6 | 0.0013 | 0.0127 |
| *Lolium perenne* | 6 | 0.0037 | 0.0371 |
| *Myosotis sp.* | 6 | 0.0044 | 0.0441 |

**Table S2. Effects of floristic richness on soil mycobiota and wheat root endospheric mycobiota diversity in the field study.** Mycobiota diversity is indicated as sequence cluster richness and Pielou’s evenness index. Random effects in the different fields were tested via linear mixed models. R^2^m is marginal R squared, denotes variance explained by just fixed effects in the model; R^2^c is conditional R squared, denotes variance explained by the entire random model; and AIC denotes Akaike's information criterion. Significant results are indicated by asterisks in brackets after Chisq value: * indicates 0.01 < P < 0.05; ** indicates 0.001 < P < 0.01, *** indicates P < 0.001. n.s: not significant. Significant results (P < 0.05) are highlighted in bold. Upward arrows denote positive effects of explanatory variables in all the models.

|  | **Soil mycobiota** | | | | **Wheat root endospheric mycobiota** | | | |
| --- | --- | --- | --- | --- | --- | --- | --- | --- |
|  | **Chisq** | **R^2^m** | **R^2^c** | **AIC** | **Chisq** | **R^2^m** | **R^2^c** | **AIC** |
| **Richness** |  |  |  |  |  |  |  |  |
| Global fungi | 0.04 (n. s) | 0.01 | 0.30 | 541 | **4.70 (_*_) ↑** | 0.10 | 0.62 | 566 |
| Ascomycota | 0.02 (n. s) | 0.01 | 0.03 | 477 | **6.70 (_**_) ↑** | 0.12 | 0.58 | 476 |
| Basidiomycota | 0.23 (n. s) | 0.01 | 0.03 | 396 | 2.00 (n. s) | 0.04 | 0.57 | 421 |
| Chytridiomycota | 0.03 (n. s) | 0.01 | 0.05 | 363 | 2.61 (n. s) | 0.06 | 0.38 | 335 |
| Glomeromycota | 0.20 (n. s) | 0.01 | 0.03 | 326 | **4.31 (_*_) ↑** | 0.07 | 0.08 | 267 |
| Zygomycota | 0.18 (n. s) | 0.02 | 0.03 | 368 | **4.98 (_*_) ↑** | 0.09 | 0.36 | 360 |
| **Evenness** |  |  |  |  |  |  |  |  |
| Global fungi | 1.08 n. s) | 0.02 | 0.26 | -181 | 0.10 (n. s) | 0.01 | 0.12 | -97 |
| Ascomycota | 0.52 (n. s) | 0.01 | 0.22 | -198 | 0.51 (n. s) | 0.01 | 0.26 | -147 |
| Basidiomycota | 2.27 (n. s) | 0.04 | 0.06 | -77 | **5.22 (_*_) ↑** | 0.10 | 0.30 | -67 |
| Chytridiomycota | 1.41 (n. s) | 0.02 | 0.04 | -72 | 1.16 (n. s) | 0.02 | 0.07 | -40 |
| Glomeromycota | 0.65 (n. s) | 0.01 | 0.12 | -73 | 2.79 (n. s) | 0.05 | 0.06 | 36 |
| Zygomycota | 0.19 (n. s) | 0.01 | 0.06 | -106 | 0.01 (n. s) | 0.01 | 0.01 | -76 |

**Table S3. Effect of weed plant identity on wheat and weed root endospheric mycobiota sequence cluster richness, number of shared sequence clusters, percentage of shared sequence clusters, wheat and weed plant aboveground weight in the controlled experiment.** Linear models were used to detect the effects of weed identity on each parameter. R^2^ denotes variance explained by the model; and AIC denotes Akaike's information criterion. Significant results are indicated by asterisks in brackets after Chisq value: * indicates 0.01 < P < 0.05; ** indicates 0.001 < P < 0.01, *** indicates P < 0.001. n.s.: not significant. Significant results (P < 0.05) are highlighted in bold.

| **Parameters** | **Chisq** | **R^2^** | **AIC** |
| --- | --- | --- | --- |
| Wheat root mycobiota sequence cluster richness | **19.51 (_*_)** | 0.17 | 741 |
| Weed root mycobiota sequence cluster richness | **83.33 (_***_)** | 0.50 | 652 |
| Number of shared sequence clusters | **23.12 (_**_)** | 0.23 | 536 |
| Percentage of shared sequence clusters | **39.27 (_***_)** | 0.33 | -119 |
| Wheat aboveground weight | **25.93 (_**_)** | 0.24 | 11 |
| Weed aboveground weight | **255.71 (_***_)** | 0.75 | -1 |
